# A cross-species rescue by mating method to interrogate gene essentiality across the *Saccharomyces* genus

**DOI:** 10.1101/2025.06.02.657496

**Authors:** Taylor K. Wang, Abigail Keller, Omar Kunjo, Maitreya J Dunham

## Abstract

The field of budding yeast research has long been empowered by the vast array of genetic tools and community resources available. Rescue by mating is one useful tool that entails mating of meiotic spores directly post-germination. This minimal cell division mating process facilitates mating based screens studying the effects of otherwise haploid lethal gene deletions. In this study, we describe the successful application of rescue by mating across two different *Saccharomyces* species: *S. cerevisiae* and *S. uvarum*. This novel inter-species tool enables studies on the evolution of essential genes within the broader *Saccharomyces* genus.

## Description

Performing genetic experiments with mutations in essential genes requires special approaches that avoid passing through lethal intermediates. Such techniques include the use of conditional mutations, transient rescue plasmids, and hypomorphic alleles. One lesser known but potentially useful approach is rescue by mating, in which a lethal mutation in a haploid is temporarily supplemented by inheritance of protein from a heterozygous diploid progenitor and can mate before death. Rescue by mating was described previously in a study elucidating the mechanisms of spore number control and spore mating in *Saccharomyces cerevisiae*, but references to the phenomenon go even further back (Taxis et al. 2005; Cox 1971).

Capable of growing as a haploid or diploid, *S. cerevisiae* often exist in the wild as diploids due to a variety of reasons (Mortimer 2000; Peter et al. 2018; Goddard, Godfray, and Burt 2005; Herskowitz 1988). Upon encountering starvation conditions, diploid yeast will trigger sporulation (Freese, Chu, and Freese 1982), during which the cell undergoes meiosis to produce four haploid spores within a protective double membrane called the ascus (Esposito and Esposito 1969). Before germination, the four post-meiotic spores within the ascus remain dormant in nutrient poor conditions. After encountering rich media, spores enter the germination cell program and begin dividing (Palleroni 1961).

The expectation for a spore which has inherited an essential gene deletion is cell death upon growth (Hartwell, Culotti, and Reid 1970; Giaever et al. 2002; Winzeler et al. 1999). Working with diploid *S. cerevisiae* strains heterozygous for an essential gene deletion, Taxis et al. discovered that spores which inherited the essential gene deletion from the mother cell were in fact capable of mating immediately after germination (Taxis et al. 2005). Spores with essential gene deletions can often undergo several cell divisions and form a microcolony before cells become incapable of further mitotic growth and undergo cell death (Rodriguez-Peña et al. 1998; Liu et al. 2015). This growth before cell death is understood to stem from the presence of sufficient maternal factors within the spore (Haarer et al. 2011) which are capable of supporting germination and limited growth before cell death. Placement of a suitable mating partner next to the germinating cell can thus result in mating and generation of a viable diploid that is heterozygous for the essential gene of interest.

Rescue by mating represents both interesting biology as well as a valuable tool. It has enabled work such as mass-screening studies of complex haploinsufficiency that relies on mating haploid essential gene deletion strains (Haarer et al. 2011). However, thus far work in the field has focused on applying this technique exclusively to *S. cerevisiae* strains. *S. cerevisiae* is only a single member of the greater *Saccharomyces* genus, which contains multiple species representing varying evolutionary distances (Borneman and Pretorius 2015). The greater *Saccharomyces* genus is of interest due to phenotypic diversity and the adaptability of *S. cerevisiae* genomic tools to other *Saccharomyces* species (Caudy et al. 2013; Bleuven et al. 2019). In addition, the mating competent *Saccharomyces* species are all capable of forming cross-species hybrids (Alix et al. 2017; Belloch et al. 2008; Bellon et al. 2013; 2015; Bernardes, Stelkens, and Greig 2017; da Silva et al. 2015). In this study we attempted to perform rescue by mating between haploid essential gene deletion *S. cerevisiae* strains and a wt *S. uvarum* mating partner—one of two most distantly related *Saccharomyces* species to *S. cerevisiae* [Figure 1A].

**Figure 1.**
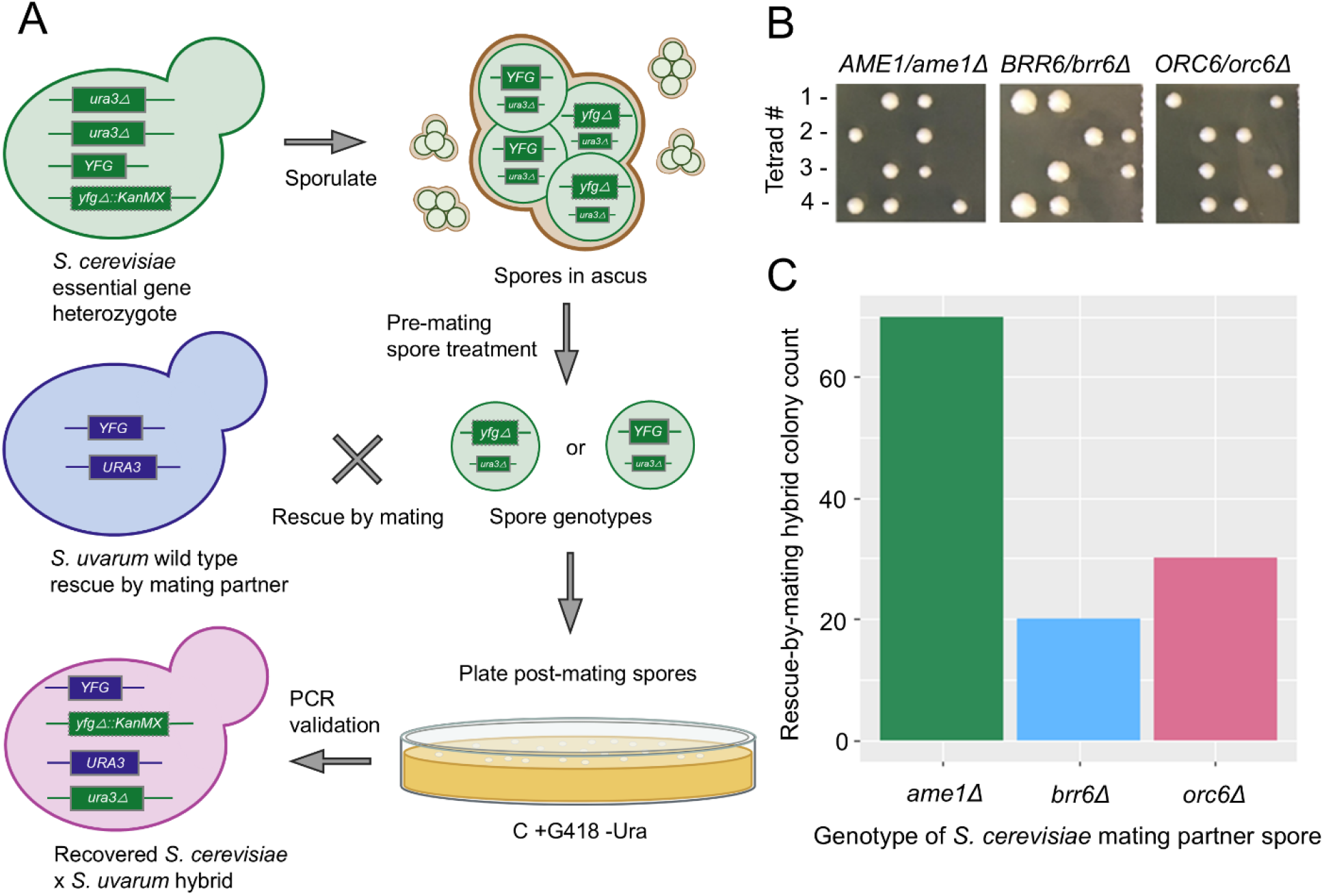
(A) Schematic depicting the *S. cerevisiae* and *S. uvarum* parent strain genotypes and the rescue by mating protocol steps. (B) Results of tetrad dissection for each of the three heterozygous knockout parent strains proving gene essentiality for *AME1, BRR6*, and *ORC6 via* 2:0 viability segregation pattern. Segregation of inviability with KanMX cassette on selection was also confirmed. (C) Recorded number of PCR and Sanger sequence verified *S. cerevisiae* x *S. uvarum* hybrid colonies recovered from rescue from mating protocol for each candidate gene.

Studying the comparative biology of *S. cerevisiae* and *S. uvarum* is of particular interest. Previous work from our lab unveiled a significant number of orthologous genes which are essential only in *S. cerevisiae*, or only *S. uvarum* (Sanchez et al. 2019). We leveraged rescue by mating as a tool to study candidate genes which are essential only in *S. cerevisiae*. The differentially essential genes studied to date can functionally complement across species. Thus, a heterozygous essential gene knockout *S. cerevisiae* strain could theoretically be used to generate gene knockout spores that could mate with a wild type *S. uvarum* partner within the rescue by mating protocol. The *S. uvarum* ortholog of the *S. cerevisiae* essential gene of interest will rescue the lethality of the essential gene knockout upon spore mating. The differentially essential candidate genes studied here include *AME1, BRR6*, and *ORC6* (Sanchez et al. 2019). The identities of these specific strains were verified by PCR and sequencing of the barcodes from the original deletion collection construction (Shoemaker et al. 1996). The essentiality of each candidate was confirmed via tetrad dissection of each respective heterozygous knockout parental strain. [Figure 1B].

We followed previous mass haploid progeny recovery methods developed to selectively destroy unsporulated diploid cells in a sporulation culture [Figure 1A] (Samsonova et al. 1985). These treatments also result in the removal of the ascus surrounding the spores. In brief, sporulation cultures of heterozygous essential gene knockouts were treated with an array of chemical and mechanical stresses as described in the Methods. The post treatment spores were split into two pools. One pool was mixed with a *MATa* ‘wild type’ *S. uvarum* partner, while the other was mated to a *MATα S. uvarum* partner. Performing rescue by mating with both mating types maximizes the number of potential rescue by mating events since both mating types of *S. cerevisiae* spores containing the essential gene knockout will have a mating partner available. These mating cultures were incubated in a roller drum overnight and surviving cells were plated onto +G418 -ura media to select for diploid hybrids and against unmated haploids: a *KanMX* cassette was present at the essential gene locus from the *S. cerevisiae* segregant while the *S. uvarum* partners were sensitive to G418, and a functional *URA3* was only present in the *S. uvarum* mating partner. Colonies on selection plates were verified as hybrids via PCR and Sanger sequencing. Final counts of recovered hybrid colonies for each mating were recorded [Figure 1C]. A small number of colonies were recovered with the correct genetic markers but that did not pass PCR validation due to the presence of both the deletion and wt *S. cerevisiae* alleles; these false positives appeared to be caused by *S. cerevisiae* segregants with aneuploidy for the chromosome carrying the essential gene locus. We found that rescue by mating between *S. cerevisiae* and *S. uvarum* partners was possible, despite previously observed low inter-species mating frequencies.

Previous work has suggested the ability of *Saccharomyces* spores to exhibit mating partner preference both within and between species (Jacobs, Gorman, and Lew 2022; Strauss et al. 2024; McClure et al. 2018; Murphy and Zeyl 2012; Murphy et al. 2006; Knop 2006; Jackson and Hartwell 1990). These studies are supported by work in our group showing the increased time for mating necessary for inter-species *S. cerevisiae* x *S. uvarum* matings compared to intra-species matings for either species: about 1 hour for intraspecies and 3 hours for interspecies. The dynamics of this mating partner preference is just one example of the questions that can be evaluated with the tool of rescue by mating across species. This novel finding that rescue by mating can result in inter-species hybrids suggests interesting biology at the level of conservation between germination and spore mating machinery across the two species within the *Saccharomyces* genus and provides another important tool in the arsenal to ask evolutionary questions across related *Saccharomyces* species.

## Materials and Methods

### Strains

All three heterozygous essential knockout *S. cerevisiae* strains were sourced from the heterozygous deletion collection originally constructed by Winzeler et al. as described there, which then had the SGA marker suite engineered later (Tong et al. 2001). Strain identity was confirmed by PCR and Sanger sequencing of the uptag barcodes. The *S. uvarum* rescue mating partners were both derived from the CBS7001 strain background.

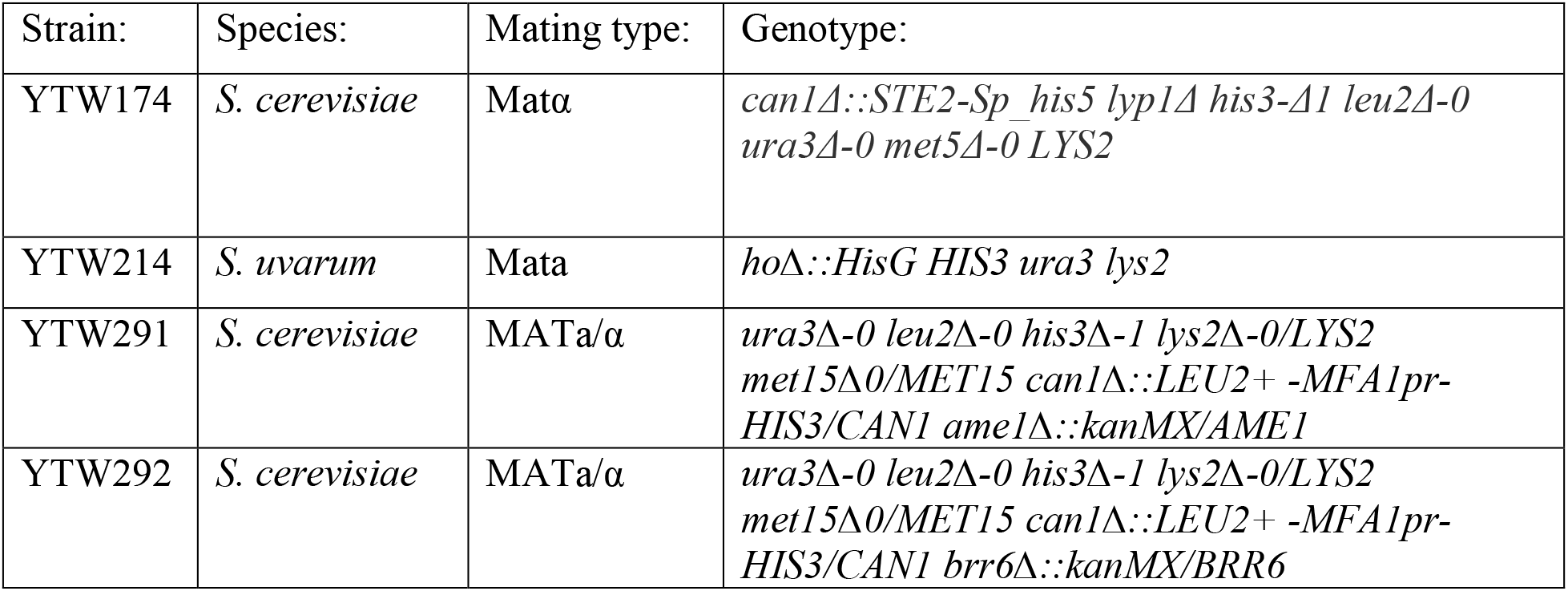
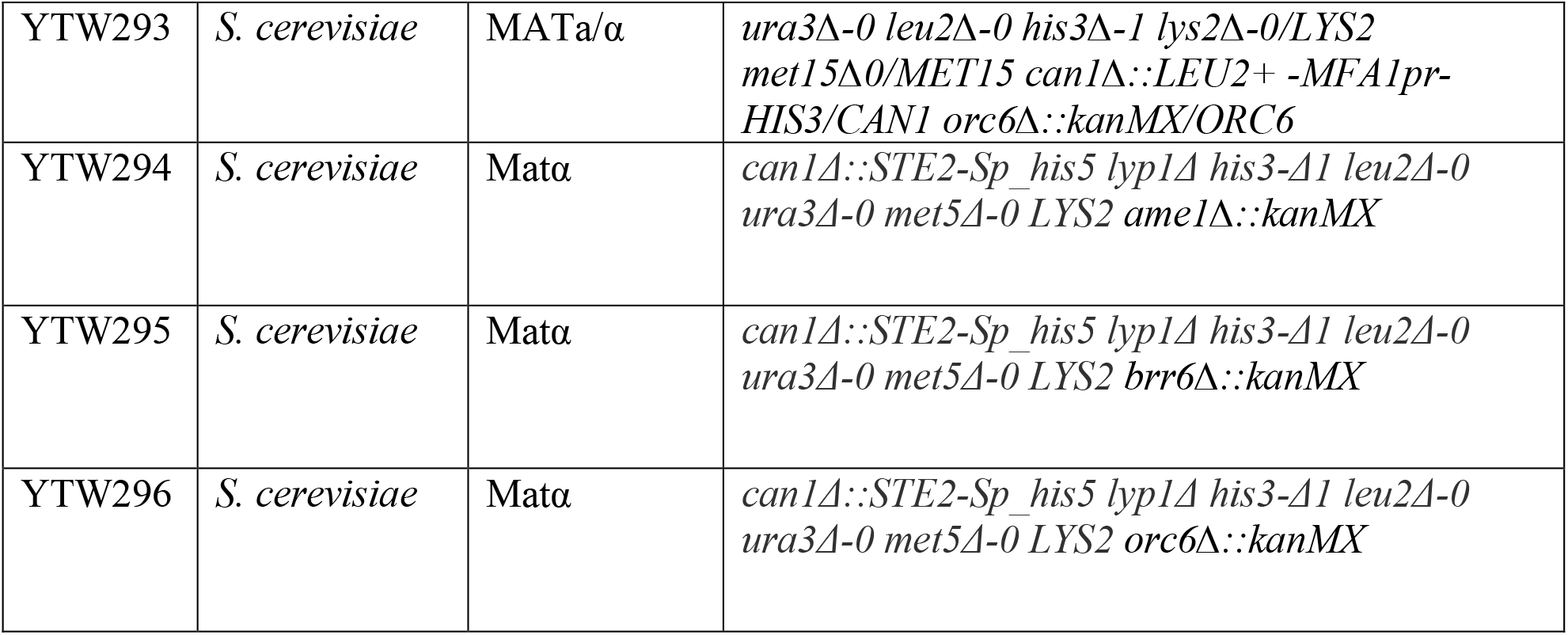

### Sporulation

A single colony of each heterozygous essential gene knockout *S. cerevisiae* strain was inoculated overnight in 3 mLs of YPD. The next day, cultures were back diluted to 0.2 OD in fresh YPD and outgrown for two more hours. 1 mL of each culture was spun down and washed twice with 1 mL of ddH_2_0. These washed cultures were each resuspended in 5 mL sporulation media consisting of 1% potassium acetate, 0.05% dextrose, and 0.1% yeast extract and placed in a roller drum at 30°C for 4-7 days.

### Rescue by mating

1 mL of sporulation culture was centrifuged for 10 seconds at 13,000xg. The pellet was resuspended in 5 mL of ddH_2_0. To this suspension 100 µL of 100 Unit/mL YLE, and 10 µL of 2- mercaptoethanol were added. Cultures were incubated overnight at 30°C with gentle shaking. The next day 5 mL of 1.5% triton X-100 was added and tubes were then incubated on ice for 15 minutes. Sonication was performed with the microtip function of a MISONIX Ultrasonic Liquid Processor. The sonicator probe was inserted into the 10 mL of spores and sonication performed as followed: microtip power setting Amplitude of 30, with three cycles total of 20 seconds of sonication and 1 minute on ice. Sonicated spores were then spun down for 10 minutes at 1200xg. The pellet was resuspended in 5 mL of 1.5% triton X-100 and vortexed vigorously. These spores were then spun down for 10 minutes at 1200xg and washed once more. After final centrifugation the spores were resuspended in 5 mL of ddH_2_0 and a 10-fold dilution counted by hemacytometer to confirm the absence of any asci or spore clumps. Spores were diluted if necessary to 1 × 10^3^ spores/mL. Matings with the *S. uvarum* rescue partner were carried out by mixing 200 µL of spores with 200 µL of log phase *S. cerevisiae* haploid mating partner at equivalent cell density. 200 µL of YPD media was added to this mating mixture, and incubated in a roller drum overnight at 30°C. The following day dilutions of the mating mixtures were plated on C -Ura +G418

## Author Contributions

Taylor Wang: Investigation, Data curation, Methodology, Formal analysis, Writing – 1^st^ draft Abigail Keller: Investigation, Data curation, Methodology, Writing – review Omar Kunjo: Data curation, Methodology, Writing – review Maitreya Dunham: Conceptualization, Investigation, Funding acquisition, Project administration, Supervision, Writing – review and editing.

